# DIMR, a Yeast-Based Synthetic Reporter System for Probing Oligomeric Transcription Factor DNA Binding

**DOI:** 10.1101/720268

**Authors:** Zachary A. Myers, Swadhin Swain, Shannan Bialek, Samuel Keltner, Ben F. Holt

**Affiliations:** Department of Microbiology and Plant Biology, University of Oklahoma, Norman, OK, USA

## Abstract

Transcription factors (TFs) are fundamental components of biological regulation, facilitating the basal and differential gene expression necessary for life. TFs exert transcriptional regulation through interactions with both DNA and other TFs, ultimately influencing the action of RNA polymerase at a genomic locus. Current approaches are proficient at identification of binding site requirements for individual TFs, but few methods have been adapted to study oligomeric TF complexes. Further, many approaches that have been turned toward understanding DNA binding of TF complexes, such as electrophoretic mobility shift assays, require protein purification steps that can be burdensome or scope-limiting when considering more exhaustive experimental design. In order to address these shortfalls and to facilitate a more streamlined approach to understanding DNA binding by TF complexes, we developed the DIMR (<u>**D**</u>ynamic, **I**nterdependent TF binding <u>**M**</u>olecular <u>**R**</u>eporter) system, a modular, yeast-based synthetic transcriptional activity reporter. As a proof of concept, we focused on the NUCLEAR FACTOR-Y (NF-Y) family of obligate heterotrimeric TFs in *Arabidopsis thaliana*. The DIMR system was able to reproduce the strict DNA-binding requirements of an experimentally validated NF-Y^A2/B2/C3^ complex with high fidelity, including recapitulation of previously characterized mutations in subunits that either break NF-Y complex interactions or are directly involved in DNA binding. The DIMR system is a novel, powerful, and easy-to-use approach to address questions regarding the binding of oligomeric TFs to DNA.

**One sentence summary:** The DIMR system provides an accessible and easy-to-use platform to elucidate DNA binding and transcriptional regulatory capacity of oligomeric transcription factor complexes

## Introduction

Transcription factors (TFs) are a fundamental component of biological control, facilitating differential gene expression in response to environmental stimuli and making complex life possible. The mechanism through which differential gene expression manifests is inherently complex, requiring coordination of stimulus perception, transcriptional and translational responses, and alteration of relevant protein activity (Crick 1970). The transcription-level integration of stimuli is of particular interest, as this process supports context-specific recruitment of RNA polymerase to a specific genomic locus (Diamond et al. 1990). This integration is accomplished through the independent, competitive, and cooperative functions of TFs and the effects of these relationships on DNA-binding and RNA polymerase recruitment (Amoutzias et al. 2008; Lickwar et al. 2012). A better understanding of the mechanisms through which transcriptional activity are modulated, particularly how TF interactions influence regulatory capacity, could facilitate the design of more precise and predictable molecular tools to address long-standing issues in the fields of agriculture, health care, and manufacturing, among others (Khalil and Collins 2010).

Current approaches to identify TF-DNA interactions are effective at *de novo* identification of binding site preferences of individual TFs *in vitro* and at identifying global binding patterns of individual TFs *in vivo*. In many cases, a combination of approaches can be used for a more complete understanding of TF targeting. In particular, many researchers have found great success in combining approaches that effectively identify binding specificity (such as Protein Binding Microarrays (PBMs), 1-Hybrid-based library screening, or DNA Affinity Purification (DAP-seq)) and lower-resolution, global-scale binding site identification (such as Chromatin Immunoprecipitation (ChIP-seq) or Assay for Transposase-Accessible Chromatin (ATAC-seq)).

While we will not focus on the general strengths of each approach or combination of approaches (see reviews, (Mahony and Pugh 2015; Jayaram et al. 2016)), one aspect of TF function that remains less explored is the impact that TF oligomerization exerts on DNA binding (Jolma et al. 2015). Most approaches, including those introduced above, have focused on understanding some aspect of DNA-binding of an assumed individual TF; however, a much more complex interplay of TF function exists than can be readily and methodically addressed with current techniques. Transcriptional regulation through multi-component TF complexes is a common theme, but our understanding of how these complexes identify and interact with specific genomic loci remains less explored (Jolma et al. 2015; Morgunova and Taipale 2017). For example, a recent study identified dramatic shifts in cognate binding sequences of bZIP homodimers compared to heterodimers (Rodriguez-Martinez et al. 2017); however, the experimental approach used in this study required purification of a large suite of individual transcription factors. An approach enabling rapid and straight-forward testing of complex TF units, with the ability to make targeted mutations in any component of the TF complex or its potential binding site, would create a new lens through which we could explore more nuanced aspects of TF-DNA interactions.

Traditional definitions of transcription factors often highlight the presence of both a DNA-binding and a transcriptional-regulation domain; however, many TFs function as obligate protein complexes, where individual components contain partial DNA-binding or transcriptional-regulation domains that are only fully-reconstituted within the context of a functional complex. One example of such an arrangement can be found in the NUCLEAR FACTOR-Y (NF-Y) transcription factors, a complex that functions as an obligate hetero-trimer of three distinct subunits, NF-YA, NF-YB, and NF-YC, and physically interacts with the DNA sequence *CCAAT*. Although this complex was initially identified over 30 years ago (Dorn et al. 1987; Jones et al. 1985), questions remain regarding the molecular forces driving specificity of the NF-Y complex to *CCAAT*. Several key observations and studies have led to persistent interest in the NF-Y, particularly in plant lineages, including: (1) the ubiquitous nature of the *CCAAT* box and, subsequently, NF-Y regulated genes, (2) the expansion of the NF-Y subunit families in plants compared to animals, where animals encode 1-2 members of each subunit, while higher plants often encode 10+ of each subunit (Siefers et al. 2009), and (3) the identification and molecular characterization of the first non-canonical NF-Y complex in plants, which replaces an NF-YA subunit for a plant-specific CCT (<u>C</u>ONSTANS, <u>C</u>ONSTANS-LIKE, and <u>T</u>OC1) domain-containing protein, leading to altered DNA binding properties (Gnesutta et al. 2017).

Published crystal structures of various NF-Y complexes, particularly of the full *Homo sapiens* NF-Y complex bound to DNA (Nardini et al. 2013), have raised increasingly complex questions regarding the specificity of the NF-Y complex for a particular *CCAAT* box. While all three NF-Y subunits have been shown to make physical contact with DNA, it appears that only NF-YA makes sequence-specific contact, inserting directly into the minor groove of the *CCAAT* box, while NF-YB and NF-YC make contacts with the DNA backbone, appropriately positioning both the DNA and NF-YA subunit for tight physical interaction (Nardini et al. 2013). Despite this understanding, the NF-Y complex appears to show more selectivity than has been experimentally derived. For example, most *CCAAT* boxes in human cell lines are not consistently bound by NF-Y complexes (Encode Project Consortium et al. 2007; Zambelli and Pavesi 2017), meaning that the mere presence of a *CCAAT* box is not predictive of NF-Y binding. Factors such as chromatin accessibility and repressive epigenetic modifications regularly limit the binding landscape of a given TF (John et al. 2011); however, the NF-Y are thought to function as ‘pioneer’ TFs that are able to bind less-accessible DNA and promote further TF complex formation (Tao et al. 2017; Oldfield et al. 2014; Donati et al. 2008). Alternatively, while the only strict requirement on NF-Y binding is the presence of the pentanucleotide *CCAAT*, the flanking nucleotides could serve as a fine-tuning mechanism for complex binding and stability. Ultimately, the source of this *CCAAT* box selectivity is not well understood and is made significantly more complex when considering the expanded family size and combinatorial complexity of plant NF-Y subunits.

In particular, we hypothesized that different NF-YB/NF-YC dimers might contribute to *CCAAT* box selectivity through interactions with the nucleotides flanking the *CCAAT* pentamer; however, comparing all possible NF-YB/NF-YC combinations with even a single NF-YA component would be a significant undertaking, with 100 possible combinations in *Arabidopsis thaliana* (Petroni et al. 2012) and 208 possible combinations in *Oryza sativa* (rice, (Hwang et al. 2016)). *In vitro* approaches requiring protein purification, such as Electrophoretic Mobility Shift Assay (EMSA), become time- and cost-prohibitive at this scale. More importantly, many proteins are recalcitrant to protein purification, and can only be isolated if appropriately truncated or co-expressed with other factors. Despite these technical limitations, EMSA analysis is well-suited for these types of investigations, and any approach intended to address similar questions would need to match or complement its capabilities. A reporter system with a large dynamic range and high sensitivity would allow scientists to address mutations that have much more modest effects on TF-DNA interactions, and could facilitate examination of more precise or biologically-relevant questions.

To address these issues and to better facilitate these types of research, we designed the DIMR (<u>**D**</u>ynamic, **I**nterdependent TF binding <u>**M**</u>olecular <u>**R**</u>eporter) system (Figure 1), a modular yeast-based transcriptional activity reporter system developed through repeated application of the synthetic biology approach of build-test-learn. The DIMR system allows for dose-dependent induction of a suite of transcription factors (effectors) and subsequent monitoring of transcriptional regulation. We validated our approach by testing the DNA binding and transcriptional activation capabilities of the Arabidopsis NF-Y^A2/B2/C3^ complex and found that the DIMR system was able to faithfully recapitulate both wild-type *CCAAT* box binding and previously-described mutations impacting DNA binding and complex formation.

**Figure 1.**
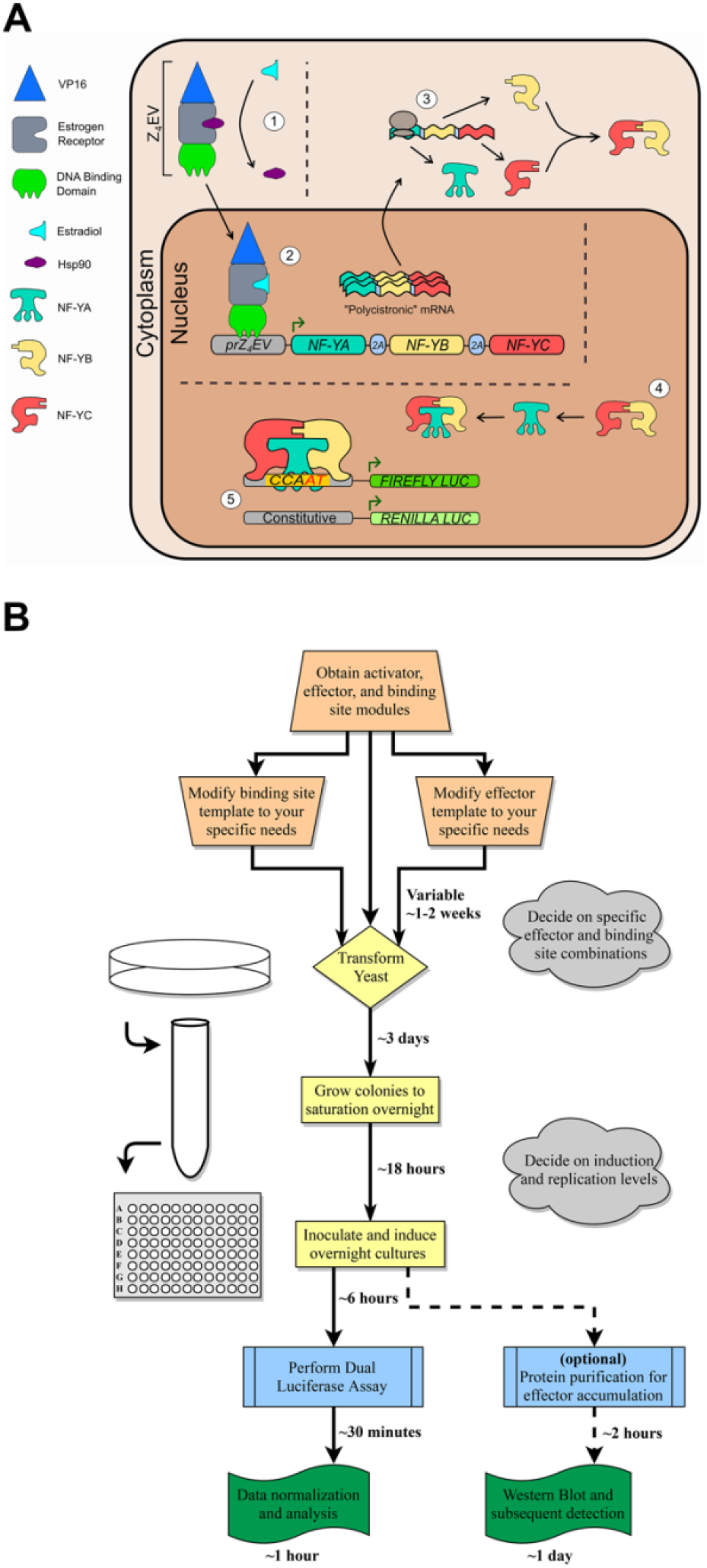
Model illustrating fundamental concepts and practical application of the DIMR system. **(A)** Circled numbers follow the flow of DIMR system activation: (1) β -estradiol activates the Z_4_EV artificial transcription factor by competing with and replacing Hsp90, allowing for nuclear accumulation of Z_4_EV; (2) Z_4_EV induces transcription of the effector cassette; (3) translation of the effector cassette mRNA produces individual effector components through 2A-mediated translational cleavage; (4) effectors translocate to the nucleus and form functional complexes; and (5) transcriptional activation is measured as the ratio of conditionally-expressed firefly luciferase activity and constitutively-expressed Renilla luciferase activities from the reporter module. **(B)** Practical application of the DIMR system, including approximate timelines for design and implementation of an individual experiment.

## Results

### DIMR system components, design philosophy, and functional description

The DIMR system is composed of three modules: (1) an **Activator Module**, encoding an inducible artificial transcription factor (ATF), (2) an **Effector Module**, encoding a single ATF-driven cassette containing viral 2A ‘cleavage’ sites between effector components, and (3) a **Reporter Module**, containing a dual-luciferase reporter composed of a constitutively transcribed *Renilla* luciferase and a conditionally transcribed Firefly luciferase (Figure 1A). Each module is carried on a yeast shuttle vector with different auxotrophic selection, and each has been designed to be as modular as possible, with restriction enzyme sites strategically placed to facilitate swapping individual effector or reporter components with minimal laboratory effort. As currently implemented, point mutations in existing effectors can be easily accomplished through commercially available site-directed mutagenesis kits, novel effector modules can be synthesized *de novo* at affordable rates, and permutations of binding sites can be cloned in as little as 3 days at a relatively low cost and with very minimal active time, resulting in a streamlined, quick, and hands-off assay to investigate TF-DNA interactions (Figure 1B). Finally, because of the synthetic biology-based approach taken during the design and implementation of the system, each component of the modules can easily be further iterated upon for improved function or to accomplish different goals.

The Activator module (Figure S1) utilizes the chimeric Z_4_EV artificial transcription factor, which contains three functional domains: (1) an engineered zinc finger with specificity to a DNA sequence not found in the genome of *Saccharomyces cerevisiae*, (2) an estrogen receptor that modulates activity and localization, and (3) a VP16 activation domain (McIsaac et al. 2013). Z_4_EV is constitutively expressed under the *ACT1* promoter (Flagfeldt et al. 2009) but remains inactive and restricted to the cytosol. Activation of Z_4_EV with the hormone β-estradiol initiates translocation to the nucleus and activation of the Effector cassette. This activation was previously shown through transcriptome profiling to induce remarkably few unintended effects, either through the action of β-estradiol itself or through off-target binding of Z_4_EV (McIsaac et al. 2013).

The Effector module (Figure S2) is designed to emulate a polycistronic message through the incorporation of viral 2A ‘self-cleaving’ sequences. Our initial designs included cassettes expressing both 2 and 3 genes, with a translationally-fused flexible linker and unique epitope tag on each (Sabourin et al. 2007). The linkers between components A – B and B – C also encode previously-validated T2A peptide sequences (Beekwilder et al. 2014), facilitating translation of individual proteins from a single mRNA. Restriction enzyme recognition sites were embedded into the coding sequences of the T2A linkers through the introduction of silent mutations. This allows easy swapping of effectors through either (1) the inclusion of the appropriate flanking linker sequence through gene synthesis (recommended, along with codon optimization (Kotula and Curtis 1991; Gustafsson et al. 2004)) or (2) PCR amplification to produce the appropriate over-hangs.

The Reporter module (Figure S3) encodes a dual luciferase reporter system, including a *Renilla* luciferase variant constitutively expressed under the *ACT1* promoter, and a conditionally expressed Firefly luciferase variant. Design of the Firefly luciferase promoter drew heavily on previously characterized yeast promoters bound by yeast NF-Y complexes. Specifically, we chose native, experimentally validated promoters of *Saccharomyces cerevisiae* whose activation required the presence of a specific *CCAAT* box (implying direct regulation by the NF-Y), and replaced the required *CCAAT* box binding site with the various permutations described below. Cloning of binding site permutations is quick and simple, requiring only a pair of appropriately designed, 5’-phosphorylated oligonucleotides and simple restriction enzyme digestion and ligation reactions.

### Validation of effector induction and cleavage

As a starting point, we tested our ability to induce accumulation of Arabidopsis NF-Y^A2/B2/C3^ effectors through the activation of Z_4_EV. After a 6-hour induction period, we were able to visualize accumulation of NF-YA2:HA (Figure 2A), NF-YB2:MYC (Figure 2B), and NF-YC3:FLAG (Figure 2C). Differences were seen in the level of protein accumulation across samples, and several instances of failed T2A-mediated cleavage were clear; however, we were able to clearly and consistently observe individual NF-Y subunits in repeated experiments.

**Figure 2.**
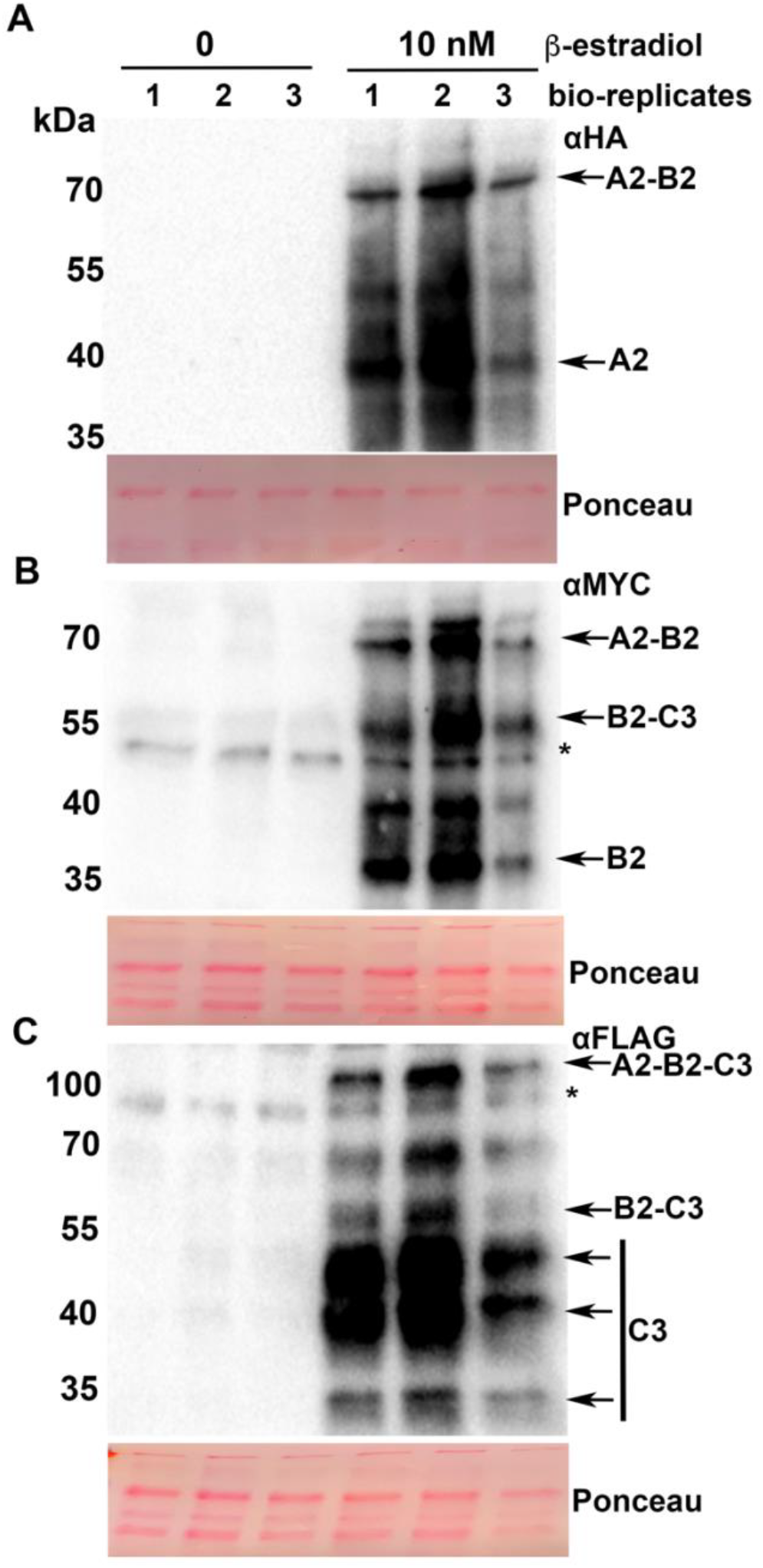
Induction and cleavage of NF-YA2, NF-YB2 and NF-YC3 effectors in yeast. Western blot analysis of **(A)** NF-YA2:HA, **(B)** NF-YB2:MYC, and **(C)** NF-YC3:FLAG accumulation in response to 10 nM β-estradiol induction for 6 hours. Three independent biological replicates, corresponding to lanes 1, 2, and 3, were tested for protein accumulation using anti-HA, anti-Myc and anti-Flag antibodies. Ponceau S staining of the PVDF membrane prior to transfer was used to test loading control. The experiment was repeated three times with similar result.

Further experiments to explore the inducibility of the effector cassette identified detectable levels of individual effector proteins with around 5 nM β-estradiol (Figure S4), with no obvious differences in accumulation between the different NF-Y subunits. A short time-course of effector accumulation at 10 nM β-estradiol identified fairly stable accumulation of individual effector proteins at 3 and 6 hours, though we often observed a peak accumulation at 6 hours and a drop-off in signal when extending to 9 hours (Figure S5). From these and other preliminary data, we established standard low- and high-induction ranges of ~1 nM and ~10 nM β -estradiol, respectively, and a standard induction length of 6 hours. While incomplete cleavage of individual effectors remains to be addressed, the induction scheme of the DIMR system is effective and robust.

### Validation of reporter activity following effector induction

Our initial system design and testing were built upon a previously-characterized NF-Y regulated *CCAAT* box from the *FLOWERING LOCUS T (FT)* promoter in Arabidopsis. This *CCAAT* box, as well as 20 flanking base pairs upstream and downstreat, was positioned immediately upstream of the minimal NF-Y regulated yeast promoter of *CITRATE SYNTHASE1 (CIT1)*. Upon induction of the NF-Y^A2/B2/C3^ effector cassette, we observed a significant, dose-dependent increase in relative luminescence (Figure 3A). Normalization of the data into a fold change value, relative to mock induction, found an average ~3.5-fold increase in high induction conditions (10 nM, Figure 3B). Despite the successful recapitulation of NF-Y DNA binding in our initial designs, we were not able to statistically distinguish between low- and high-induction conditions, and the maximum fold changes observed were lower than desired. A larger dynamic range of reporter activation would facilitate addressing more nuanced questions, such as the DNA binding impact of individual effector mutations or changes in cognate binding sites.

**Figure 3.**
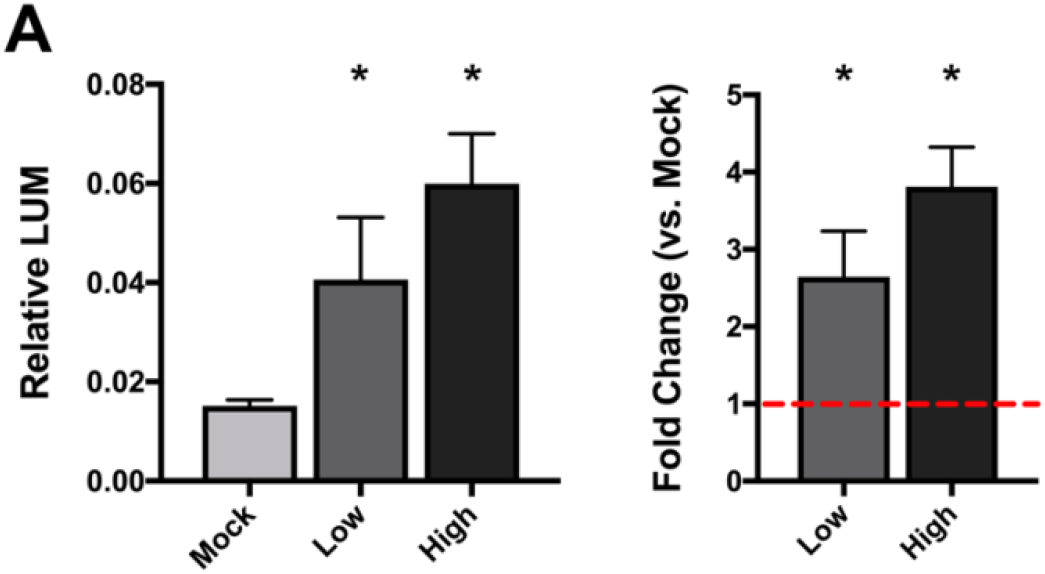
DIMR system test of NF-YA2/B2/C3 on FT CCAAT box before system optimization. **(A)** Relative luminescence of mock, low, and high concentration β-estradiol (0, 1uM, and 10uM, respectively) treated samples, and fold change levels (compared to mock) of low and high treated samples. Each condition includes at least 6 biological replicates. Asterisks indicate a significant increase between mock and treated samples, as determined through two-way ANOVA (p < 0.01). Error bars represent 95% confidence intervals.

### Refinement of binding site design

From these initial tests, we next sought to refine the DIMR system for increased reporter dynamic range. We focused our system tuning on two approaches: reducing mock-level signal and increasing maximum activation. To decrease mock-level signal, we designed and tested different NF-Y regulated minimal promoter architectures to identify reporters with lower basal level activation. We focused on promoters for the genes *CIT1* and *ASPARAGINE SYNTHETASE1 (ASN1)*, driven by the above-described *FT* CCAAT box footprint (Figure 4A). Luciferase activity levels were lower in mock-treated samples with the *ASN1-*based promoter compared to the originally-tested *CIT1*-based promoter (Figure 4B). Notably, this reduction in mock-level activation translated to a larger fold change of ~6x in *ASN1*-based reporters when comparing mock and high induction conditions.

**Figure 4.**
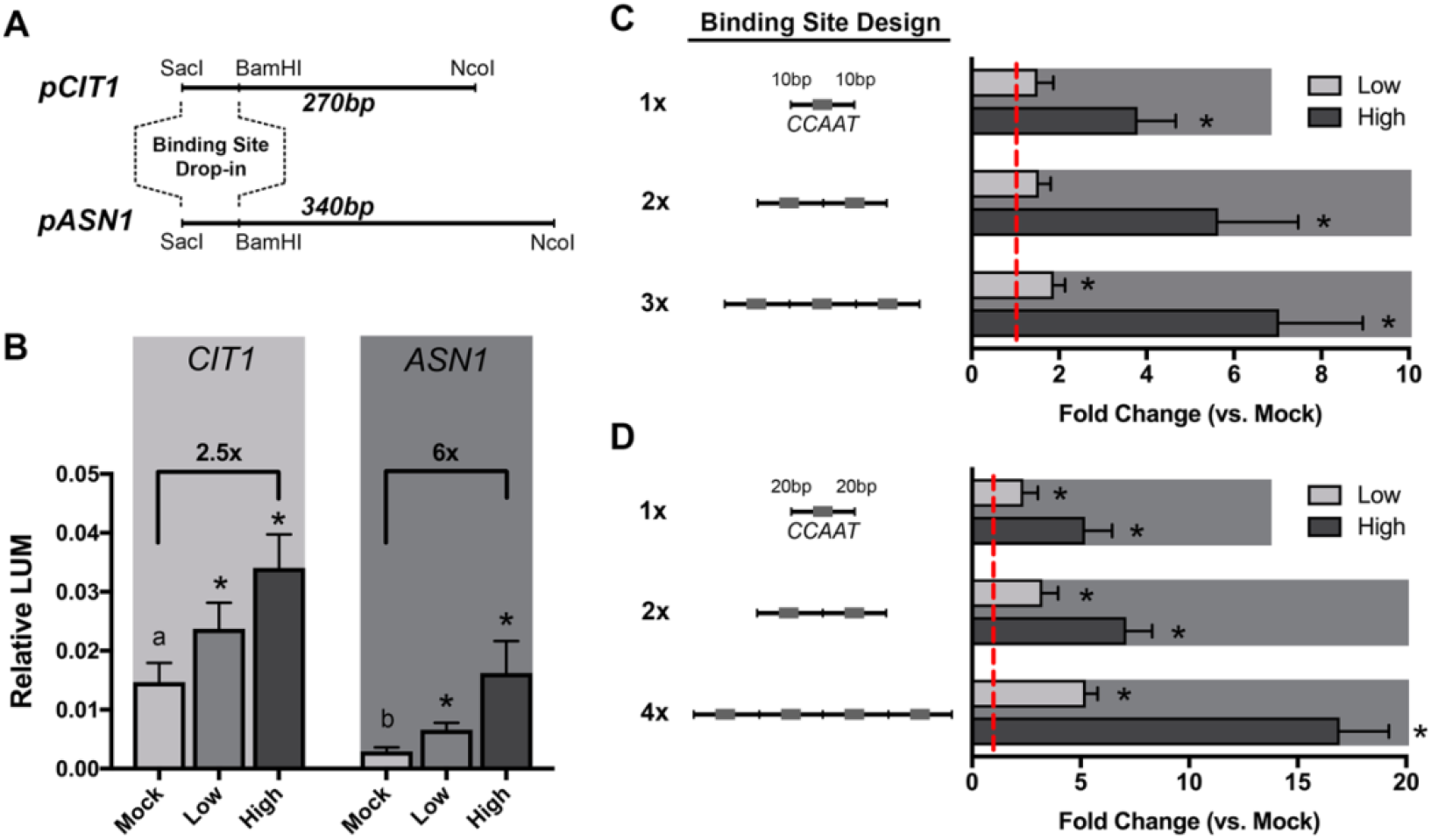
Parameter refinement of the DIMR system to optimize reporter dynamic range. **(A)** Model of two tested promoter architectures, derived from the *CIT1* and *ASN1* native yeast promoters. Binding site permutations were cloned into the reporter module through flanking SacI / BamHI restriction sites. **(B)** Effects of different promoter architectures of *CIT1* and *ASN1* on DIMR system activation of NF-Y^A2/B2/C3^ on *FT CCAAT* box. **(C, D)** Effects of binding site multimerization and altered spacing between binding sites. In **(C)**, each binding site footprint was 25bp, while in **(D)**, each footprint was 45bp. Both sets were testing binding of NF-Y^A2/B2/C3^ on *FT CCAAT* box permutations in the *ASN1*-based promoter architecture. Error bars represent 95% confidence intervals. Asterisks above individual bars indicate statistical significance over matched mock-treated samples, as determined through two-way ANOVA (p < 0.01). Asterisks above brackets connecting two bars indicate significance between the two conditions, determined similarly. Letters above mock-induced bars indicate significance groups between the different promoter architectures, also determined through two-way ANOVA (p < 0.01).

To increase our maximum signal, we additionally tested the effects of different numbers of binding sites and the spacing between them. Multimerization of available binding sites can result in an increase in observed transcriptional activity (Khalil et al. 2012), but this larger DNA footprint can be more difficult to accommodate in cloning. To abrogate this, we also tested the effects of reducing the length of each individual binding site footprint. First, we attempted to cut the *CCAAT* box footprint roughly in half by including only 10 base pairs of flanking sequences in single, double, and triple binding site configurations (Figure 4C), and found that multimerization of the binding site increased total system activation. Finally, we combined the original, larger *CCAAT* box footprint into single, double, and quadruple binding site configurations (Figure 4D). In this set, the quadruple binding site configuration stands out at a much-improved ~15-fold increase in reporter activity from mock. The larger DNA footprint of the quadruple binding site approach required a slightly modified cloning approach, where two pairs of annealed, phosphorylated oligos were simultaneously ligated into the Reporter module; however, we were able to preserve the flexibility of the DIMR system and avoided the need for further gene synthesis for individual binding site permutations.

### Validation and assessment of refined binding site setup

With an increased signal level upon system activation, we began testing the activation requirements of the DIMR system. First, we examined dose-dependent system activation levels of the NF-Y^A2/B2/C3^ complex on the quadruple *CCAAT* box over a wide range of β-estradiol induction levels (Figure 5A). This induction gradient was analyzed through 4-parameter logistic regression (4PLR), with an R^2^ value of 0.942. No system activation was observed at induction levels below 1 nM, while activation peaked between 10 and 100 nM β-estradiol. The dose-dependent activity of the system was most pronounced between 1 nM and 10 nM, with a half maximal effective concentration (EC_50_) of ~2.2 nM and total span of ~10.5-fold change. 4PLR is a widely used metric in chemistry, biochemistry, and computational biology to describe cooperative action, and in this case, statistically supports the idea that the individual NF-Y subunits are working interdependently in an induction-dependent manner to bind the *CCAAT* box.

**Figure 5.**
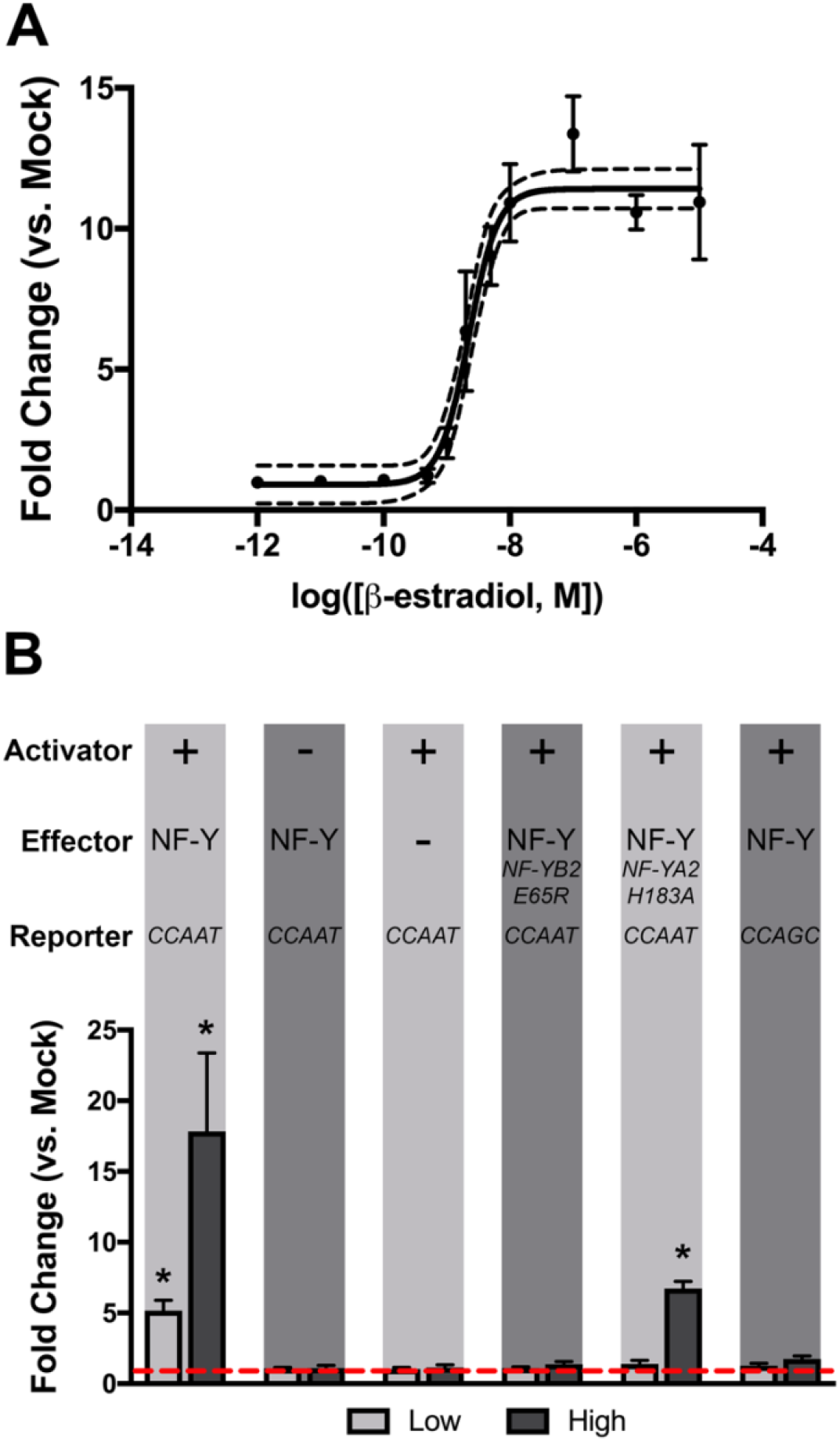
Functional assessment of DIMR system activation. **(A)** Dose response curve of NF-Y^A2/B2/C3^ on the 4x *FT CCAAT* box in the ASN1 promoter architecture. The solid line fits the dataset to a 4-parameter logistic regression, with 95% confidence intervals plotted with dashed lines (R^2^ = 0.942). **(B)** DIMR system requirements for NF-Y/CCAAT-mediated activation. Module components are generally described above each condition: +, present; -, absent; NF-Y, NF-Y^A2/B2/C3^. Error bars represent 95% confidence intervals. Asterisks above individual bars indicate statistical significance over matched mock-treated samples, as determined through two-way ANOVA (p < 0.01); all bars lacking asterisks were found to be nonsignificant compared to matched mock-treated samples.

To further solidify the correlation between effector induction and system activation, we tested the impacts of loss of individual DIMR modules, previously described mutations in NF-Y effectors, and changes in the CCAAT box (Figure 5B). Replacing either the Z_4_EV activator cassette or the NF-Y effector cassette with empty vectors resulted in a complete loss of system activation upon induction. Importantly, we found that mutation of the *CCAAT* box to *CCAGC* eliminated system activation. These three pieces of data collectively suggest that the observed induction responses are accomplished through the action of the NF-Y complex on the *CCAAT* box driving the firefly luciferase reporter.

We also performed further system tests with the *NF-YB2 E65R* point mutation, a previously characterized mutation shown to prevent interaction of the NF-YB/NF-YC dimer with NF-YA (Sinha et al. 1996; Siefers et al. 2009). As expected, this point mutation abolished system activation (Figure 5B). While many highly-conserved residues in each of the NF-Y subunits have been previously identified and characterized to completely break DNA binding (Sinha et al. 1996; Kim et al. 1996; Sinha et al. 1995), we wanted to examine novel mutations that could help reveal the evolutionary constraints acting on the NF-Y complex. To this end, we examined the effects of one particular point mutation at a residue thought to be directly involved in *CCAAT* box binding, *NF-YA2 H183A*. With the native histidine residue replaced with an alanine at this location, we still observed a moderate level of system activation (Figure 5B). This alanine-replacement approach is traditionally used to replace residues that contribute to sequence-specificity with an unobtrusive and relatively-inert alanine that is unlikely to mediate sequence-specific interactions (Luscombe et al. 2001).

Because we had previously observed instances of failed 2A site function in the form of fused effector components, we directly tested whether these fused effectors were functional by creating point mutations that break the existing 2A sites. We found that fusion of the NF-YA and NF-YB components did lower maximum reporter activity compared to a 2A-cleaved trimeric system or a fusion between NF-YB and NF-YC, but each instance of broken 2A cleavage was still able to significantly activate the reporter system (Figure S6). While not all transcription factor complexes will be able to effectively form and regulate DNA while fused together, this should be tested for each group of transcription factors being tested.

Finally, we tested whether yeast NF-Y orthologs might be able to compete with or take the place of our effector module-encoded Arabidopsis NF-Y proteins. Sequentially removing one NF-Y subunit from the otherwise-complete effector module induced no system activation (Figure S7), suggesting that native NF-Y orthologs are not responsible for any significant activation of the reporter. While interactions between NF-Y orthologs of different species has been demonstrated (Calvenzani et al. 2012), we hypothesized that the level of induction through the activator and effector modules would quickly saturate any unintended interactions. Further, not all yeast NF-Y orthologs are thought to be constitutive expressed, with at least one ortholog only expressed in response to non-fermentable carbon sources (Bourgarel et al. 1999).

### Validation of CCAAT-based NF-Y binding

To further explore the affinity of the NF-Y complex to the *CCAAT* box, we tested permutations of the *CCAAT* box with variation at the 3’ end, corresponding to all possible combinations of *CCA*<u>***NN***</u> (Figure 6). We focused on variation at these locations because the crystal structure of the DNA-bound human NF-Y complex identified many more direct interactions with NF-YA and the first three bases of the *CCAAT* box than the fourth and fifth bases. Beyond this, our earlier exploration of the *NF-YA2 H183A* mutation suggests possible differences in the way human and plant NF-Y complexes bind DNA, an idea further supported by key differences in the otherwise-conserved linker region of *NF-YA2*. This linker region was proposed to be important for precise positioning of flanking α-helices for proper NF-Y complex stabilization and DNA binding, and *NF-YA2* is the only family member in Arabidopsis with this unusually long linker region.

**Figure 6.**
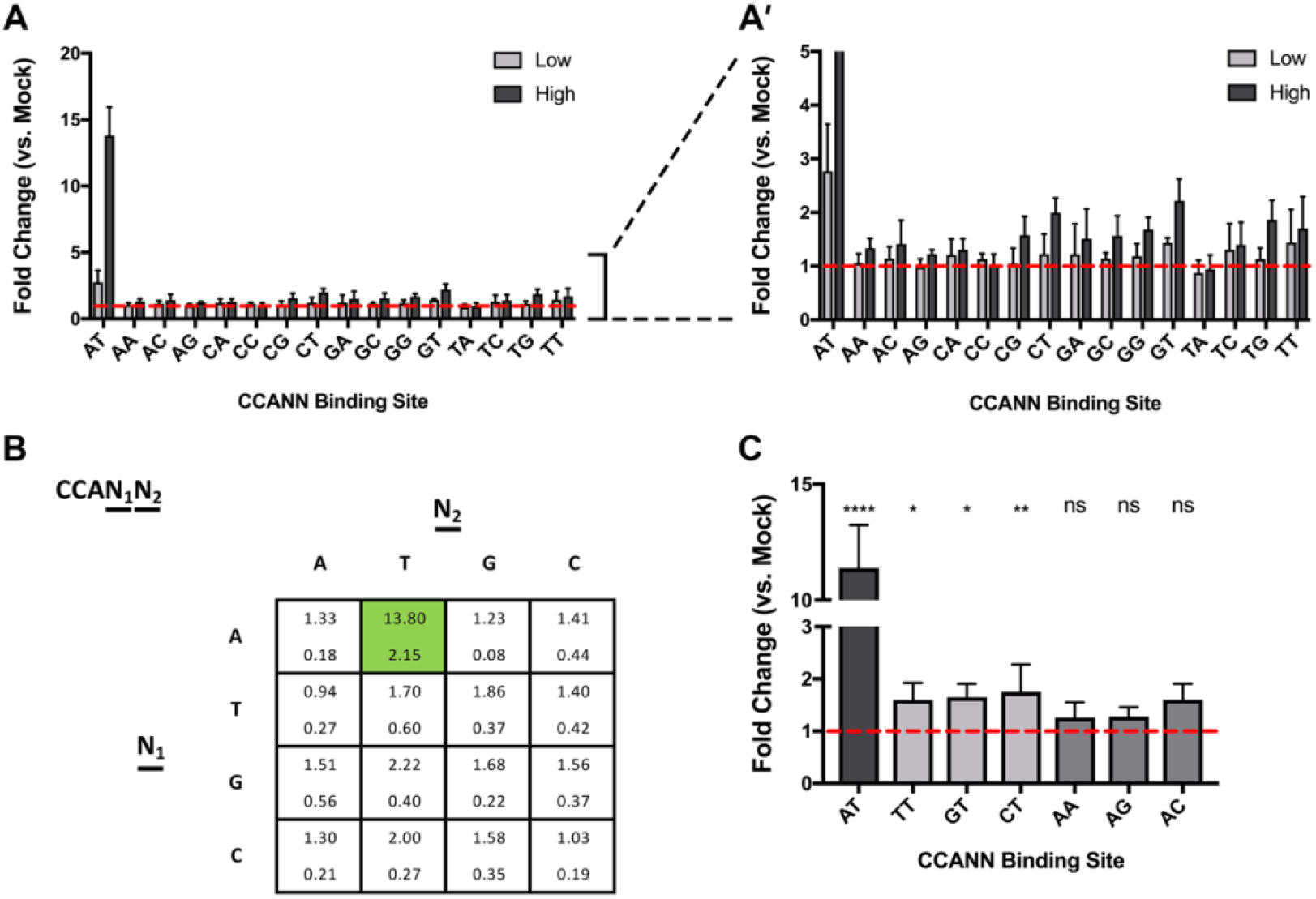
DIMR-based validation and investigation of NF-YA2/B2/C3 CCAAT binding. **(A)** System activation of NF-Y^A2/B2/C3^ on the *FT CCAAT* box, with all possible binding site permutations corresponding to *CCA*<u>***NN***</u>. Error bars represent 95% confidence intervals. Asterisks above individual bars indicate statistical significance over matched mock-treated samples, as determined through two-way ANOVA (p < 0.01); all bars lacking asterisks were found to be nonsignificant compared to matched mock-treated samples. **(B)** Graphical representation of the data presented in panel A. Each cell corresponds to a specific permutation of *CCA*<u>***NN***</u>, and contains the average fold change (top) and standard deviation (bottom). (C) A targeted comparison of CCANT and CCAAN binding site permutations. Asterisks indicate significance compared to mock induction as determined by two-way ANOVA with Sidak’s multiple comparison adjustment (****, p < 0.001; **, p < 0.01; *, p < 0.05).

Unsurprisingly, the highest-activated binding site of the *CCA*<u>***NN***</u> suite was *CCA*<u>***AT***</u> at ~14-fold over mock, with no other binding site permutation activating greater than 2.2-fold over mock (Figure 6A). Among these non-CCAAT binding sites, we saw a small increase in reporter activity in *CCA*<u>***N***</u>*T*, but not in *CCAA*<u>***N***</u> (Figure 6B). A targeted comparison of these two binding site permutations identified significant increases relative to mock in all *CCA*<u>***N***</u>*T* binding sites, but none other than *CCAAT* in the *CCAA*<u>***N***</u> variants (Figure 6C). While further investigation is necessary, this observation runs counter to what is currently understood about DNA binding by the NF-Y complex in humans, as a direct physical interaction between NF-YA has been identified at the fourth position, but not the fifth position, of the CCAAT box.

## Discussion

### Using DIMR to investigate TF complex-DNA interactions

In spite of the importance and ubiquitous nature of multi-component transcription factor assemblies, relatively few molecular or biochemical approaches have been devised to explore the intricacies of DNA binding by these multimeric complexes. Current and widely adopted methods are not well-suited for many questions, particularly those that could benefit from extensive mutagenesis-based analyses, such as alanine scanning mutagenesis to probe the relative contributions of DNA binding amino acids within a given TF. After fine-tuning and optimizing our induction and binding site schemes, we ultimately produced a sensitive, straightforward, and versatile reporter system for assaying transcription factor complex DNA interactions. As one example of this type of study, we examined the effect of permutations of the *CCAAT* box and identified a slightly higher level of system activation when altering bases at position 4 than position 5 (*CCA*<u>***N***</u>*T* vs. *CCAA*<u>***N***</u>). Our system bypasses the need for protein purification or deep sequencing, two common requirements for other approaches. The proposed system can easily be scaled to simultaneously include dozens of effector or binding site permutations with relative simplicity, and requires only relatively basic lab equipment. The injector-fitted, luminescence-based plate reader is likely the only equipment not commonly found in a standard molecular lab, and these types of equipment are often housed in core facilities available to most universities or institutes.

### Incomplete cleavage at T2A sites in effector module

Our initial designs sought to leverage viral 2A peptide sequences to generate equimolar amounts of individual effectors through a cleavage mechanism during translation, an important goal when considering obligate oligomeric TF complexes. We use the term cleavage throughout; however, strictly speaking, the end result is an inability to form the peptide bond between two adjacent amino acids, not true cleavage of an existing peptide bond (Kim et al. 2011). Because eukaryotes do not utilize the polycistronic mRNA approach so common in bacteria, researchers have turned to 2A sites and other approaches to emulate this effect. While not a perfect solution to this problem, we found that viral T2A sites worked sufficiently well for our needs here. It is important to note, however, that incomplete cleavage of the effector cassette does alter the interpretation of system activation. Therefore, with the shortcomings of this design, we can test whether a transcription factor complex can bind *a particular consensus sequence*, but we cannot directly address whether a particular complex *can or cannot form*. While this distinction is fairly minor in many cases, particularly when working with previously-described complexes, it is an important limitation of the system that should be addressed in future iterations. Unfortunately, no completely effective eukaryotic-based system has been described to recreate the polycistronic mRNA system so widely employed in bacteria (Blount et al. 2012). However, new 2A peptide sequences are still being identified and have been seen to vary in efficiency from organism to organism (Luke et al. 2010), raising the possibility that a 2A site more efficient in yeast remains to be identified.

To circumvent this issue, we have begun designing effector cassettes with individual components driven by separate inducible promoters. These separate promoters each contain the Z_4_EV -bound consensus sequence, but utilize different minimal promoters to reduce recombination within yeast (Broach et al. 1982). Whether this modification will lead to relatively-equal protein accumulation is unknown; however, it should at least provide a framework to begin further modifying, tweaking, and improving the system for our specific needs.

### Considerations and improvements on the DIMR system

While much of the parameter refinement presented above should help inform novel design considerations, some applications will require more significant alterations. In particular, issues are likely to arise where effector complexes do not possess intrinsic transcriptional activation potential or when a binding site is bound and autoactivated by endogenous yeast proteins. In cases where complexes completely lack activation potential, an activation tag might be fused to one or more effectors (Knop et al. 1999); however, whether this approach could overcome active transcriptional repression is unclear. Similarly, instances of reporter autoactivation might be addressed through fusion of one or more effector components to a transcriptional repressor domain (Edmondson et al. 1996) and measuring inducible *reductions* of reporter activity. While we have not yet worked through either of these concerns, the modular design of the system should allow for relatively straight-forward modification and testing.

### Teasing apart the details of NF-Y DNA binding: a case for DIMR

Despite consistent investigations into the mechanistic function of the NF-Y in animals over the past 30 years, many aspects of complex formation and DNA binding remain relatively unexplored in plant NF-Y orthologs. First, a significant amount of plant NF-Y research, including most DNA-binding assays, has been conducted using only conserved domains of each NF-Y subunit. In fact, the survey presented above represents one of the first comprehensive examinations of DNA binding and transcriptional regulation by a complete, full-length plant NF-Y trimeric complex. While the core domains of each NF-Y subunit show significant conservation across all eukaryotic lineages, the flanking termini show remarkable divergence from one another. The biological significance of these flanking regions has remained elusive in most cases, though several protein-protein interactions with NF-Y complexes are thought to be mediated through NF-Y terminal domains (Cao et al. 2011). A better understanding of the functions of the less-conserved termini is critical for plant NF-Y research in particular, as the vast majority of differences between individual members of expanded NF-Y gene families are found in these flanking regions. The DIMR system is capable of facilitating a systematic domain-swap approach of the flanking regions of NF-YB or NF-YC paralogs, and could potentially uncover changes in NF-Y complex affinity or specificity achieved through modulation of the specific NF-Y subunits within a functional complex.

Like *NF-YB* and *NF-YC*, the conserved domains of *NF-YA* family members are remarkably similar to one another. One critical exception to this observation is a 4-5 amino acid elongated linker sequence encoded in the *NF-YA2* subunit. This linker is positioned between two highly-conserved alpha helices that are each responsible for NF-Y complex formation or DNA binding of the mature complex, and in addition to influencing the positions of the flanking alpha helices, this linker also makes physical contact in several places with the sugar-phosphate backbone of the *CCAAT* box (Nardini et al. 2013; Romier et al. 2003). While experimental evidence is necessary to understand the impact and significance of this atypical linker, it is possible that NF-Y complexes containing NF-YA2 have slightly different DNA binding profiles or NF-Y complex dissociation constants. It should be noted, however, that NF-YA2 is likely the best-characterized plant NF-YA subunit, as it has been a major focus of research through its regulation of photoperiodic flowering. While the majority of our observations regarding NF-YA2 DNA binding and complex formation have supported the crystal structure of the DNA-bound human NF-Y complex, many important aspects of NF-Y form and function have not been thoroughly investigated and developed.

In fact, many of the most exciting and significant advances in NF-Y research have occurred in the last two years through the identification and characterization of the first non-canonical NF-Y complex. This complex utilizes the NF-YB2/NF-YC3 dimer to stabilize interactions between CONSTANS (CO) and its consensus binding site. This complex, termed NF-CO, recognizes a *CCACA* motif found in the proximal promoter of the *FLOWERING LOCUS T (FT)* gene and is critical for proper photoperiodic floral induction (Gnesutta et al. 2017). The interaction between CO and the NF-YB/NF-YC dimer is mediated through a conserved, plant-specific CCT (<u>***C***</u>*ONSTANS*, <u>***C***</u>ONSTANS-LIKE, and <u>***T***</u>OC1) domain that is found in over 40 genes in Arabidopsis (Griffiths et al. 2003; Farré and Liu 2013), raising the possibility of a significantly-expanded pool of potential NF-Y or NF-CCT complexes. Excitingly, several families of CCT domain containing proteins have been extensively studied and shown to be of critical importance for photoperiodic flowering (Griffiths et al. 2003), circadian clock entrainment and maintenance (Mizuno 2004), and seedling environmental responses (Reyes et al. 2004). Unfortunately, testing hypotheses concerned with these non-canonical NF-CCT complexes is far from straightforward, with over a decade passing between the initial suggestions that CO might function through NF-YB/NF-YC to a complete set of work describing the NF-CO complex. Systematic testing of the ability of different NF-CCT complex to form and bind DNA is a massive undertaking, with over 4,000 possible NF-CCT complexes to test. Further, while the two are clearly related, the CCT domain shows key differences from the DNA-binding domain of NF-YA, and variation within CCT members occurs at residues predicted to be important for DNA binding in aligned NF-YA sequences. A systematic mutational analysis focusing on the differences between important residues of aligned NF-YA and CCT members could create a map of critical DNA-binding residues, and inform the search for cognate binding sequences of various NF-CCT complexes.

The DIMR system is well-suited to address these types of questions on structure and function, as supported by our analysis of the *NF-YA2 H183A* point mutation and our comparison of NF-Y^A2/B2/C3^ system activation in *CCA*<u>***N***</u>*T* and *CCAA*<u>***N***</u> permutations. The *NF-YA2* histidine residue at position 183 is thought to make sequence-specific contact with the *CCAAT* box (Gnesutta et al. 2017), but reduced DIMR system activation is still observed in the *H183A* point mutation. Importantly, when aligning NF-YA and CCT DNA binding domains, this residue diverges between, but not among, many clades and sub-clades. Considering this pattern of divergence, it is intriguing to consider that this particular residue might contribute to DNA binding specificity of different NF-CCT complexes. The identification of an NF-YA mutation at this residue that retains some activity could be relevant to this observation in an evolutionary context, as a sub-optimal variant is thought to facilitate further functionalization by providing a novel evolutionary trajectory (i.e., without such a permissive intermediate, purifying selection cannot be overcome to reach a different high-fitness state (Poelwijk et al. 2007; Anderson et al. 2015)). Further, the relative tolerance of wild-type system activation at *CCA*<u>***N***</u>*T* compared to *CCAA*<u>***N***</u> is interesting to consider in light of the evolutionary constraints acting on the functionalization of NF-Y and NF-CCT complexes. This permissiveness could provide space for evolution to act, allowing suboptimal interactions to lead to novel functions of the complex. Direct testing of these types of evolutionarily significant hypotheses is technically challenging, requiring more exhaustive mutagenic approaches such as alanine scanning mutagenesis and subsequent functional validation (Cunningham and Wells 1989). The DIMR system presented here was designed to bridge the gap in tools necessary to more directly address these types of more nuanced questions.

## Acknowledgements

The authors would like to thank Dr. Scott Russell for assistance with grant administration. Additionally, we thank Dr. Daniel Jones, Dr. Marielle Hoefnagels, Dr. Laura Bartley, and Andrew Willoughby for their insightful and constructive comments on the preparation of this manuscript. Finally, we thank the National Science Foundation for funding this research through MCB EAGER 1747539.

## Methods

### Yeast strain selection

The tests presented here all utilized the BY4735 (ATCC 200897) strain (*MATα ade2Δ::hisG his3Δ200 leu2Δ0 met15Δ0 trp1Δ63 ura3Δ0*), a derivation of the S288C laboratory strain. BY4735 was obtained and is available from the American Type Culture Collection (https://www.atcc.org/).

### Yeast growth, transformation, and induction conditions

Yeast strains were uniformly grown at 30C. Wild type strains were grown with constant agitation in YAPD liquid medium and transformed through the traditional lithium-acetate based approach, as previously described. Transformants were selected after 2-3 days growth on synthetic medium lacking Leucine, Tryptophan, and Uracil (L^−^ W^−^ Ura^−^).

For DIMR system induction, colonies were first grown to saturation in liquid L^−^ W^−^ Ura^−^ media. Saturated cultures were then diluted 1:1 to a total sample volume of 500µL with fresh liquid L^−^ W^−^ Ura^−^ media containing either ethanol (mock treatments) or one of a range of β-estradiol concentrations. These 500µL induction samples were cultured on deep, 2mL 96-well plates for 6 hours before sample collection for dual luciferase reporter testing. Plates were covered with Breathe Easy strips during induction.

Unless directly stated otherwise, each system test was conducted after 6 hours of induction, with treatments corresponding to mock induction (ethanol only), low induction (1-1.5nM β-estradiol), and high induction (10nM β-estradiol).

### Module design and construction

The individual modules were each incorporated and carried on a different yeast shuttle vector – *pRS314* and *pRS315* for the effector and activator constructs, respectively, and a modified *pRSII316* removing an *NcoI* recognition site within the *URA3* coding sequence through site directed mutagenesis (described below) for the reporter construct. Initial module designs were synthesized by Biomatik^®^, while individual effector drop-in components were synthesized by Genewiz^®^.

The activator module encodes the Z_4_EV artificial transcription factor, driven by the constitutive *ACT1* promoter (Flagfeldt et al. 2009) and flanked by the *CYC1b* terminator (Curran et al. 2013). The effector module uses the Z_4_EV-bound promoter to drive expression of the ‘polycistronic’ effector mRNA. Each component has been translationally fused to a 5x-glycine flexible linker (Sabourin et al. 2007) and a unique epitope tag (component A: HA, component B: MYC, component C: FLAG). Viral T2A sites were incorporated after the HA and MYC tags to facilitate cleavage into individual effector components. The entire cassette is flanked by the CYC1b terminator. The reporter module situates the two luciferase variants in a tail-to-tail fashion. The constitutively-active Renilla luciferase is expressed under the ADH1 promoter (McIsaac et al. 2013) and is flanked by the ADH1 terminator (Curran et al. 2013), while the conditionally-expressed Firefly luciferase is driven by a modified yeast minimal promoter (*ASN1* and *CIT1* presented here, (Sundseth et al. 1997; Nevoigt et al. 2006)) that includes previously described NF-Y bound CCAAT box sequences (Cao et al. 2014; Gnesutta et al. 2017; Siriwardana et al. 2016) and is flanked by the CYC1b terminator. All plasmids used in this study are freely available, and are described in Table S1.

### Binding site and promoter architecture cloning, site directed mutagenesis

Drop-in Binding Site (DIBS) cloning was accomplished through annealing of 5’ phosphorylated oligos and subsequent ligation into the reporter module. DIBS primers were designed and ordered as pairs of complementary oligos that were then annealed together by boiling for 5 minutes in annealing buffer (10mM Tris HCl, pH 8.0, 1mM EDTA, 50mM NaCl) and slowly cooling to room temperature. The reporter module was restriction enzyme digested (SacI/BamHI, New England Biolabs) and dephosphorylated (Quick Dephosphorylation Kit, New England Biolabs), then purified and concentrated through the Zymo DNA Clean and Concentrator kit (PN). Ligations were set up with 3:1 ratios of insert:backbone free ends with ~50-150ng of purified backbone.

Site directed mutagenesis was conducted on effector entry clones through New England Biolabs Q5^®^ Site Directed Mutagenesis Kit, following manufacturer’s instructions. Primers for mutagenesis were designed with guidance of the NEBaseChanger™ tool, and mutagenesis was verified through restriction enzyme digestion (where appropriate) and Sanger sequencing through the University of Oklahoma’s Biology Core Molecular Lab.

### Western blots

Total yeast proteins were extracted using Y-PER™ Yeast Protein Extraction Reagents (Thermo Scientific, Cat no: 78991) following manufacturer’s protocol. The protein concentrations were measured using Pierce™ BCA Protein Assay Kit (Thermo Scientific, Cat no: 23225) following manufacturer’s protocol, and was quantified on a Synergy HTX Multi-Mode Reader (BioTeK, USA) at 562 nm wavelength. The protein concentrations were calculated through generation of a BSA standard curve. A total of 3 μg of protein was loaded to Mini-PROTEAN TGX Stain-Free Gels (10% gel, BIO-RAD). Proteins were transferred to standard PVDF membranes through an OWL semi-dry transfer apparatus, and the presence of NF-YA2:HA, NF-YB2:MYC, or NF-YC3:FLAG was probed with high affinity anti-HA primary antibodies (Roche, catalog no. 11 867 423 001), anti-MYC (Abcam, catalog no. ab9106), and anti-FLAG (Abcam, catalog no. F3165) followed by rabbit anti-rat (Abcam, catalog no. ab6734), goat anti-rabbit (Abcam, catalog no. ab205718), and goat anti-mouse (Abcam, catalog no. ab6789) HRP-conjugated secondary antibodies, respectively. A Bio-Rad ChemiDoc XRS imaging system was used for visualizing the protein blot after incubations with ECL plus reagent (GE Healthcare, catalog no. RPN2132).

### Dual luciferase assays

Dual Luciferase assays were performed on the BioTek^®^ Synergy™ HTX multi-mode plate reader fitted with dual reagent injectors, using the Illumination™ Firefly & Renilla Luciferase Enhanced Assay Kit (Goldbio^®^, cat# I-920) per manufacturer instructions, with the following alterations: (1) we used only 5µL of each culture, (2) samples were not spun down and/or washed, and (3) we used half-volume injections of both luciferase buffers. Our initial system tuning followed the manufacturer instructions more strictly, and we could determine either no functional difference or minor improvements between manufacturer instructions and our modified instructions (data not shown).

### Statistical approaches

Statistics were calculated through Graphpad Prism. One-way ANOVA on relative luminescence values was used for comparisons of DIMR system activation to mock levels. When denoting significance in fold change-based metrics, the relative luminescence data was used for statistical analysis.

### Figure and model construction

Individual graphs were generated through Graphpad Prism, while full figures were composed in Adobe Photoshop CC2018. The model in Figure 1A was constructed in Inkscape, while the flowchart in Figure 1B was constructed through draw.io v10.4.5 (https://draw.io/).

---

**Figure S1.**
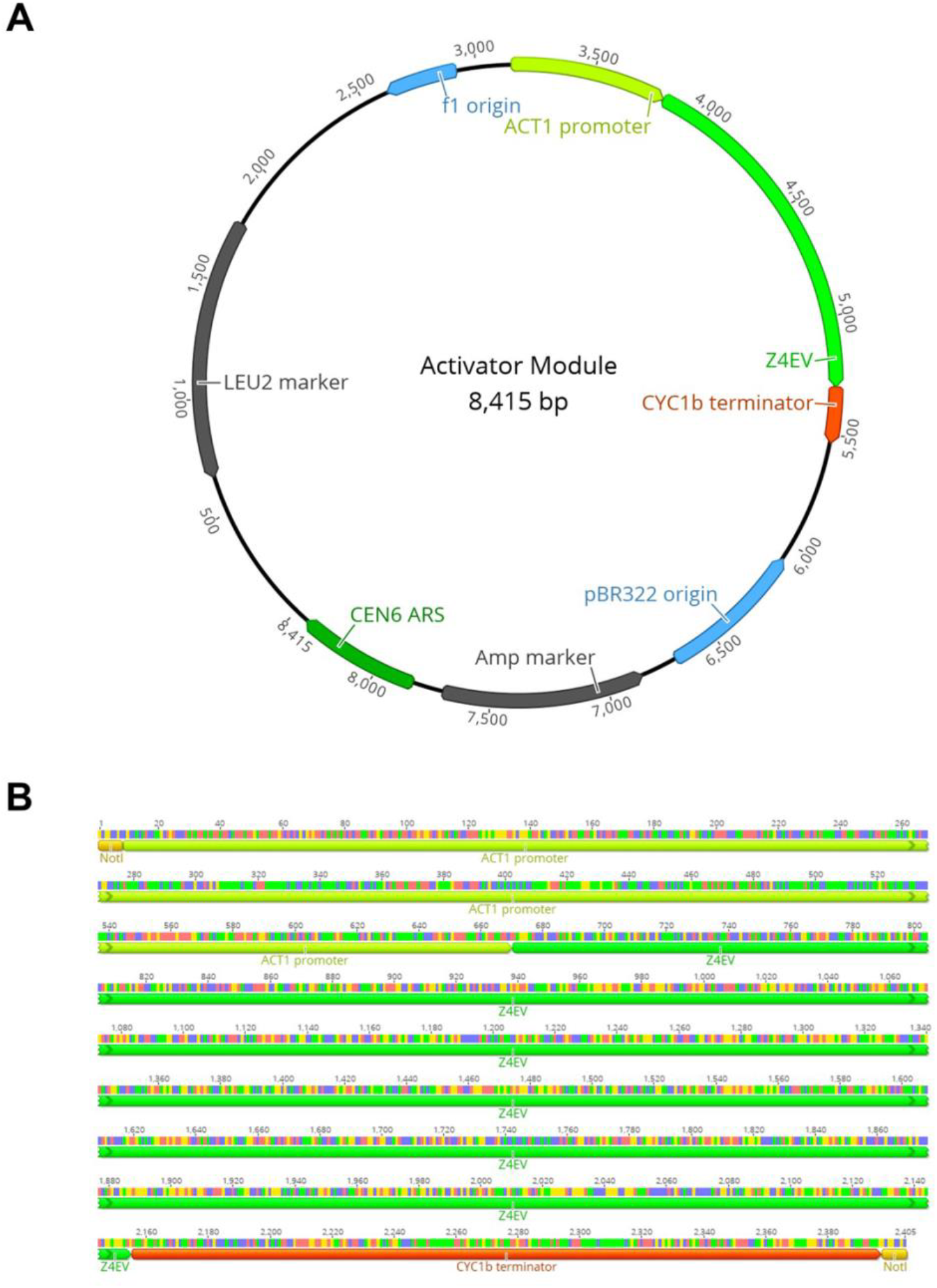
Main features of the Activator module. (A) Plasmid map of the DIMR Activator (pDIMR_A###) module showing important features and their orientation. (B) Zoomed-in view of the relevant Activator components.

**Figure S2.**
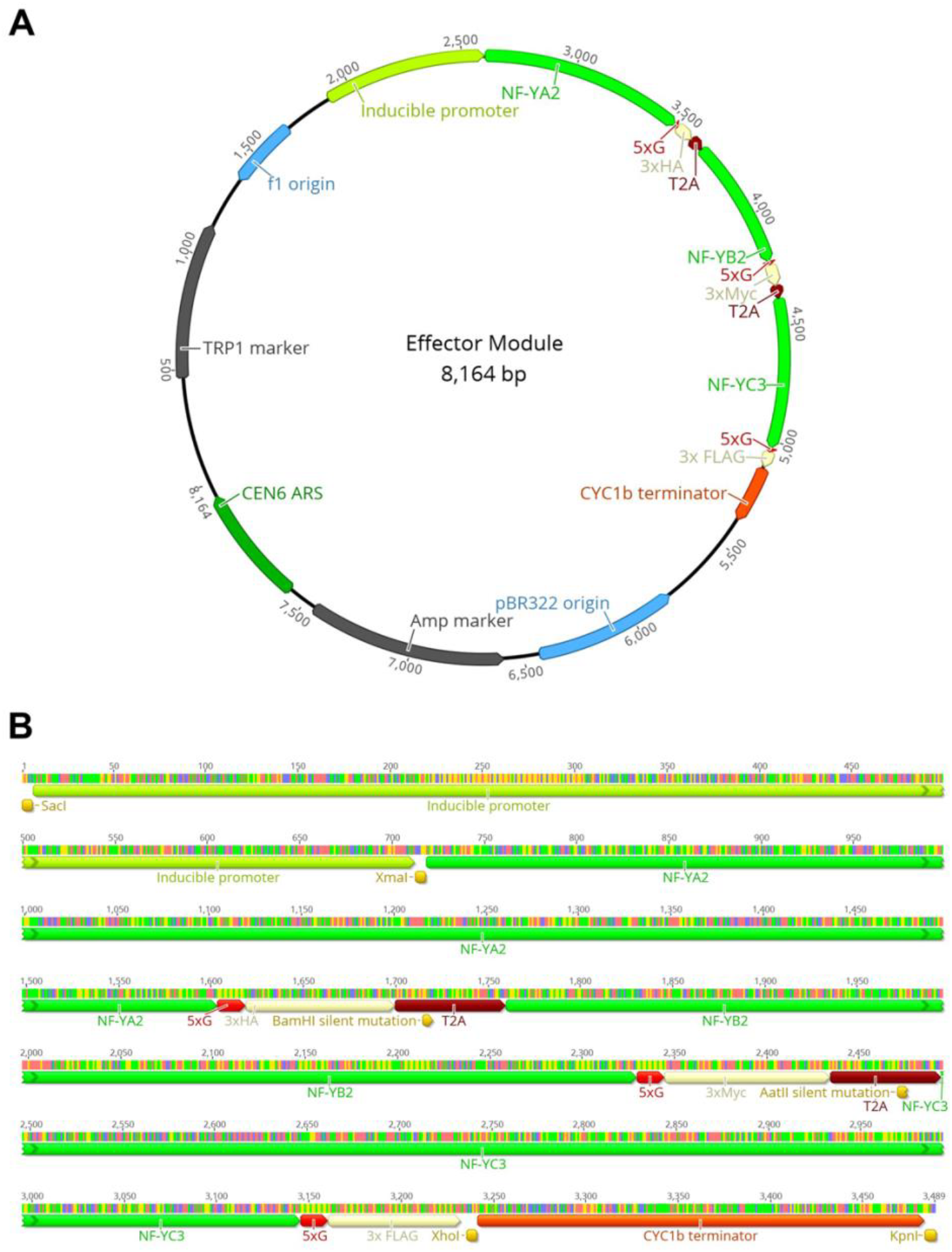
Main features of the Effector module. (A) Plasmid map of the DIMR Effector (pDIMR_E###) module showing important features and their orientation. (B) Zoomed-in view of the relevant Effector components, including important restriction enzymes used for swapping new components into or out of the system.

**Figure S3.**
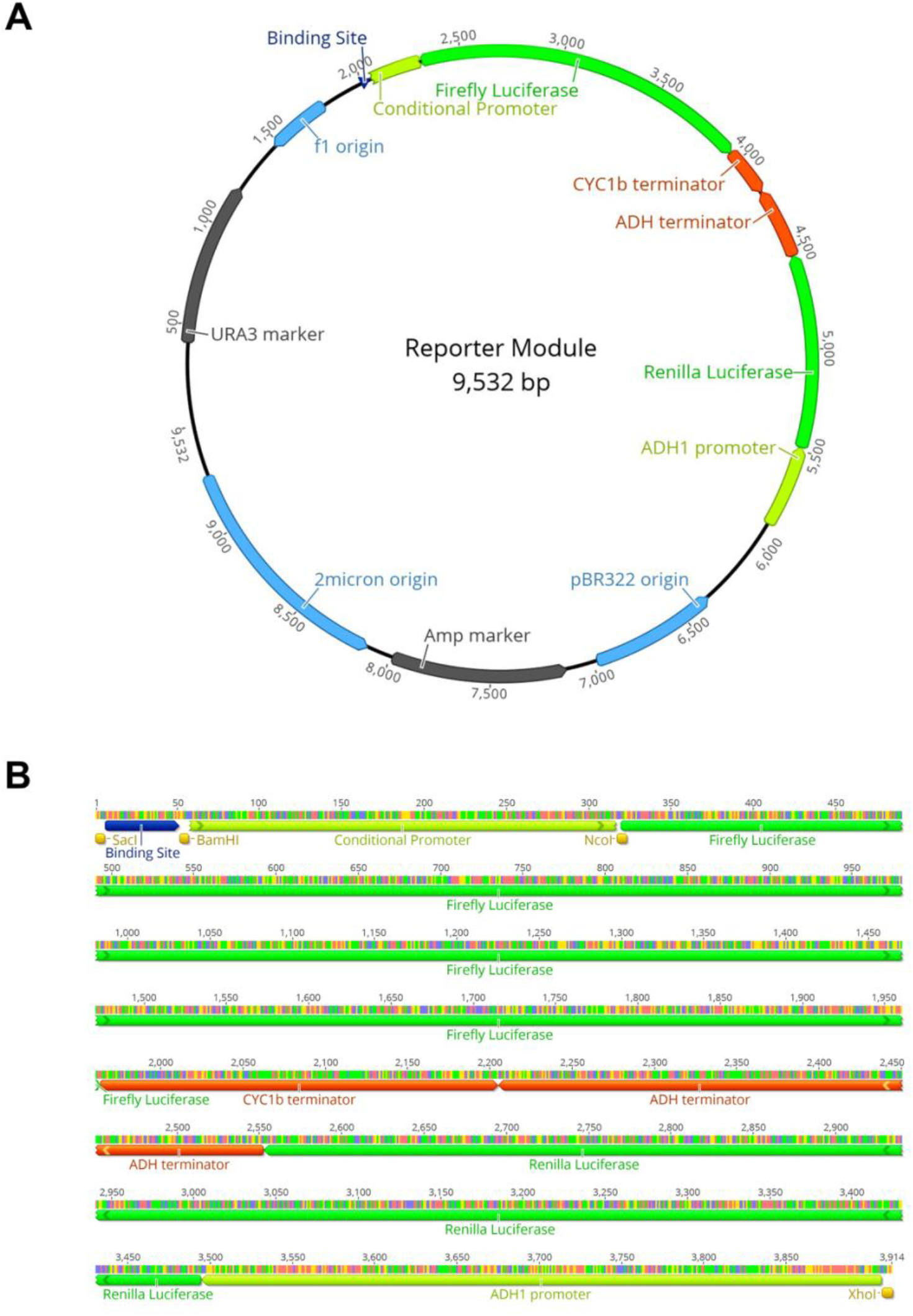
Main features of the Reporter module. (A) Plasmid map of the DIMR Reporter (pDIMR_R###) module showing important features and their orientation. (B) Zoomed-in view of the relevant Reporter components, including important restriction enzymes used for Drop-In Binding Site (DIBS) cloning.

**Figure S4.**
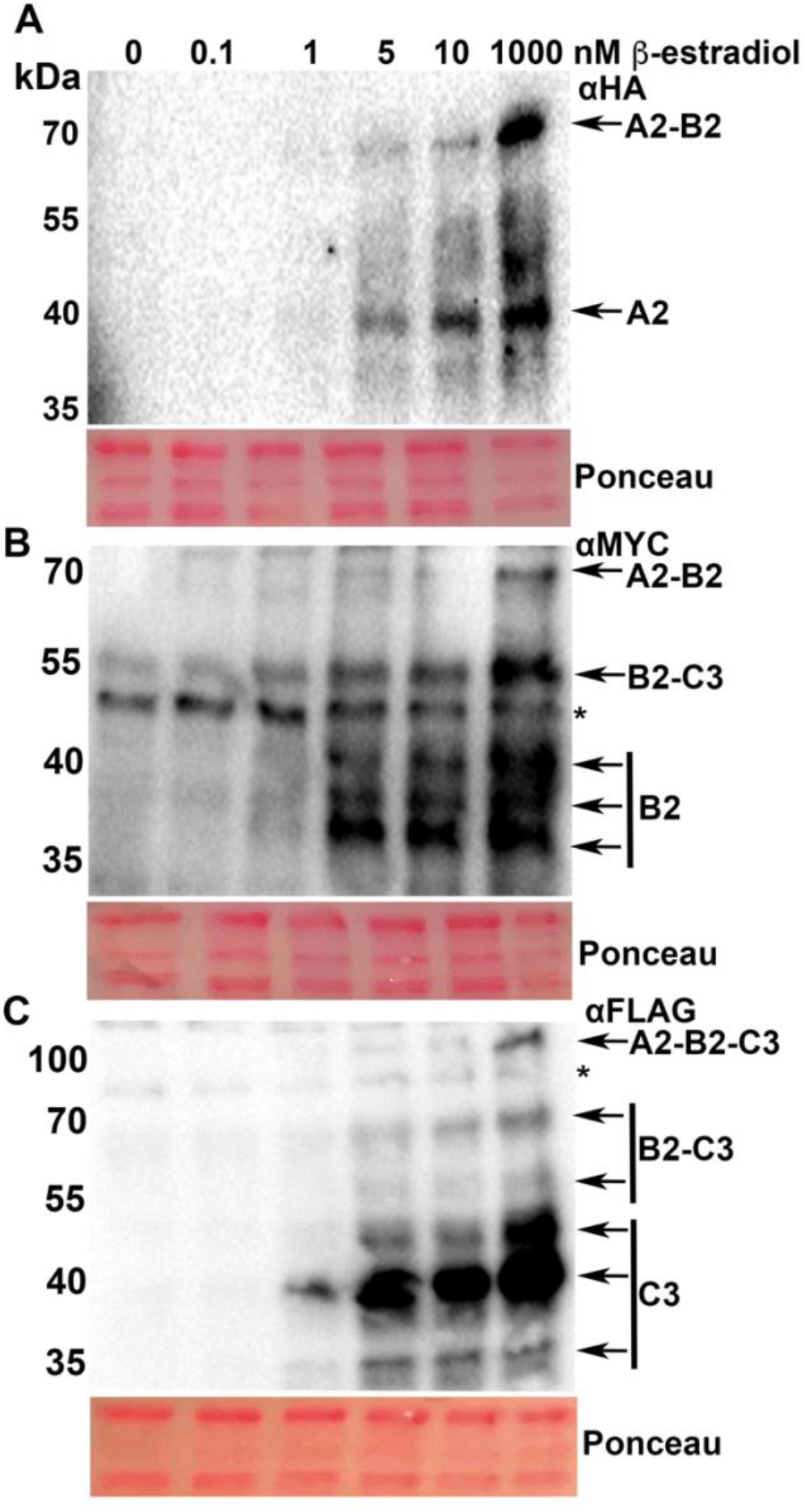
Induction gradient of NF-YA2, NF-YB2 and NF-YC3 effectors in yeast. Western blot analysis of **(A)** NF-YA2:HA, **(B)** NF-YB2:MYC, and **(C)** NF-YC3:FLAG accumulation in response to varying levels of β-estradiol induction for 6 hours. A single biological replicate was tested for protein accumulation using anti-HA, anti-Myc and anti-Flag antibodies. Ponceau S staining of the PVDF membrane prior to transfer was used to test loading control. The experiment was repeated twice with similar results.

**Figure S5.**
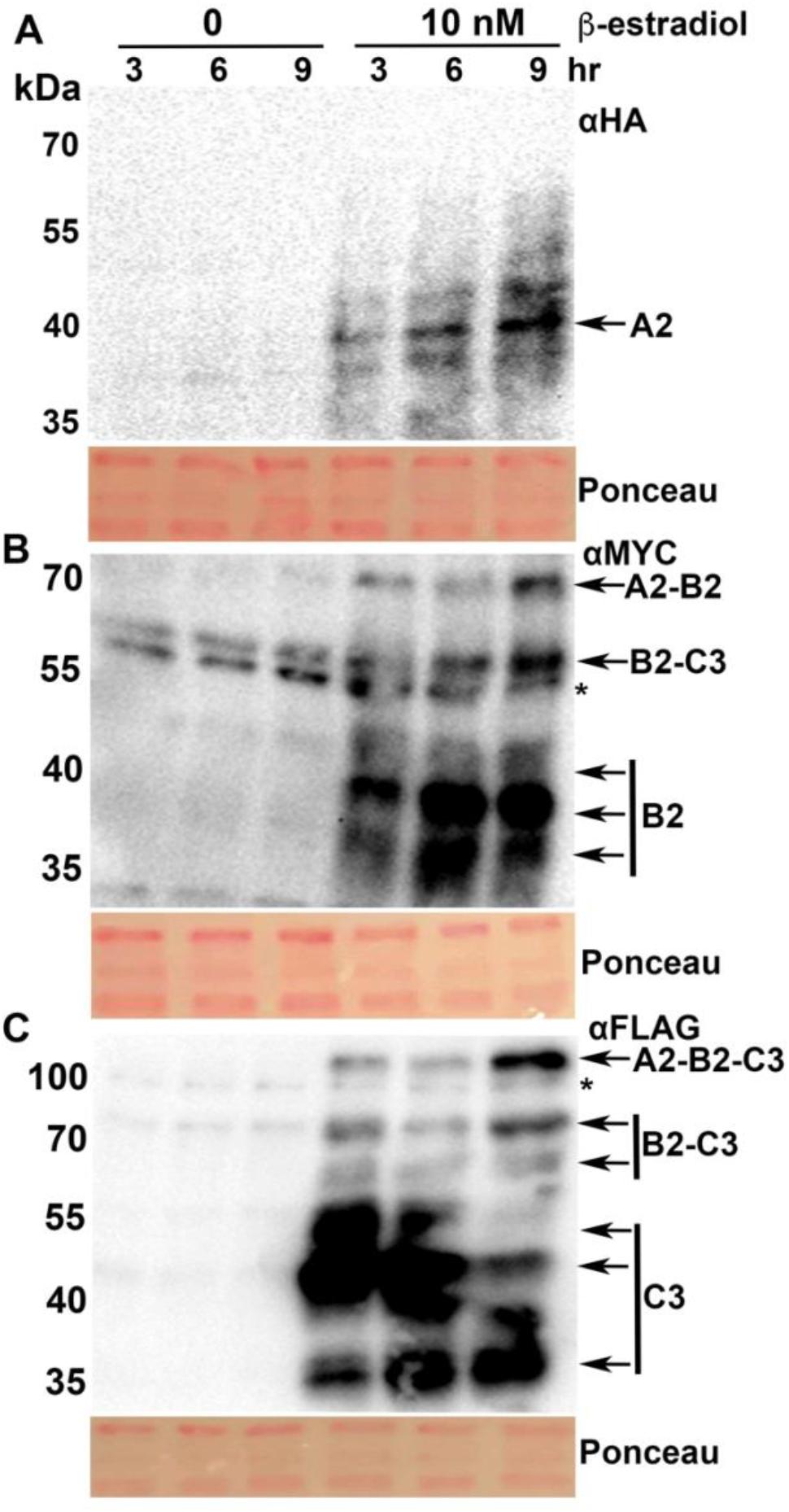
Induction time-course of NF-YA2, NF-YB2 and NF-YC3 effectors in yeast. Western blot analysis of **(A)** NF-YA2:HA, **(B)** NF-YB2:MYC, and **(C)** NF-YC3:FLAG accumulation in response to 10nMf β-estradiol induction for 3, 6, and 9 hours. A single biological replicate was tested for protein accumulation using anti-HA, anti-Myc and anti-Flag antibodies. Ponceau S staining of the PVDF membrane prior to transfer was used to test loading control. The experiment was repeated twice with similar results.

**Figure S6.**
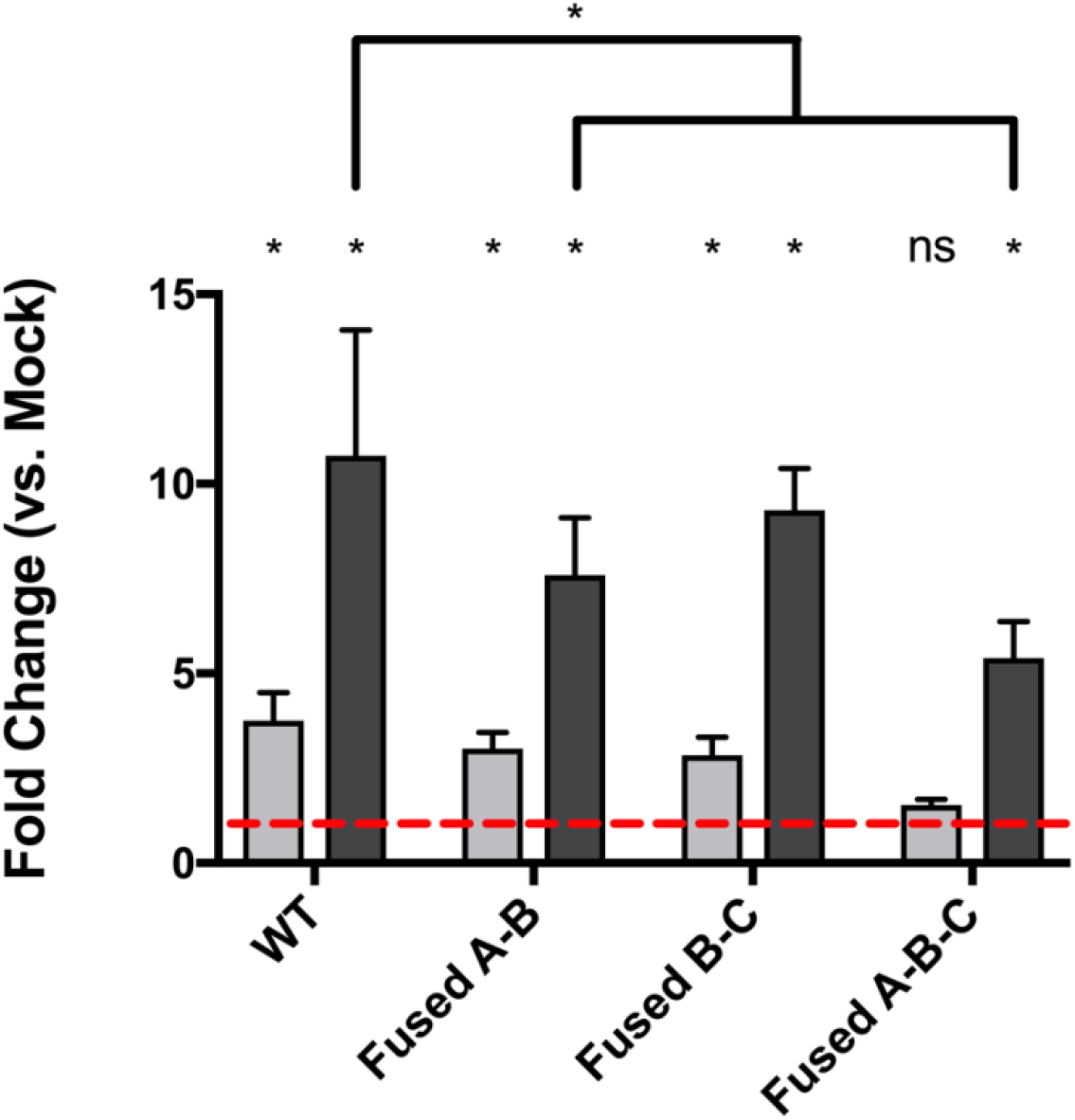
Activation of the DIMR system in effector cassettes with broken T2A sites. Point mutations were generated in the T2A sites between NF-YA / NF-YB, between NF-YB / NF-YC, and in both T2A sites simultaneously. T2A sites were broken through a proline to alanine mutation near the end of each T2A site, as previously described (Beekwilder et al. 2014). Error bars represent 95% confidence intervals. Asterisks above individual bars indicate statistical significance over matched mock-treated samples, as determined through two-way ANOVA (p < 0.01). Brackets with asterisks indicate significance between induced samples. ns, not significantly different.

**Figure S7.**
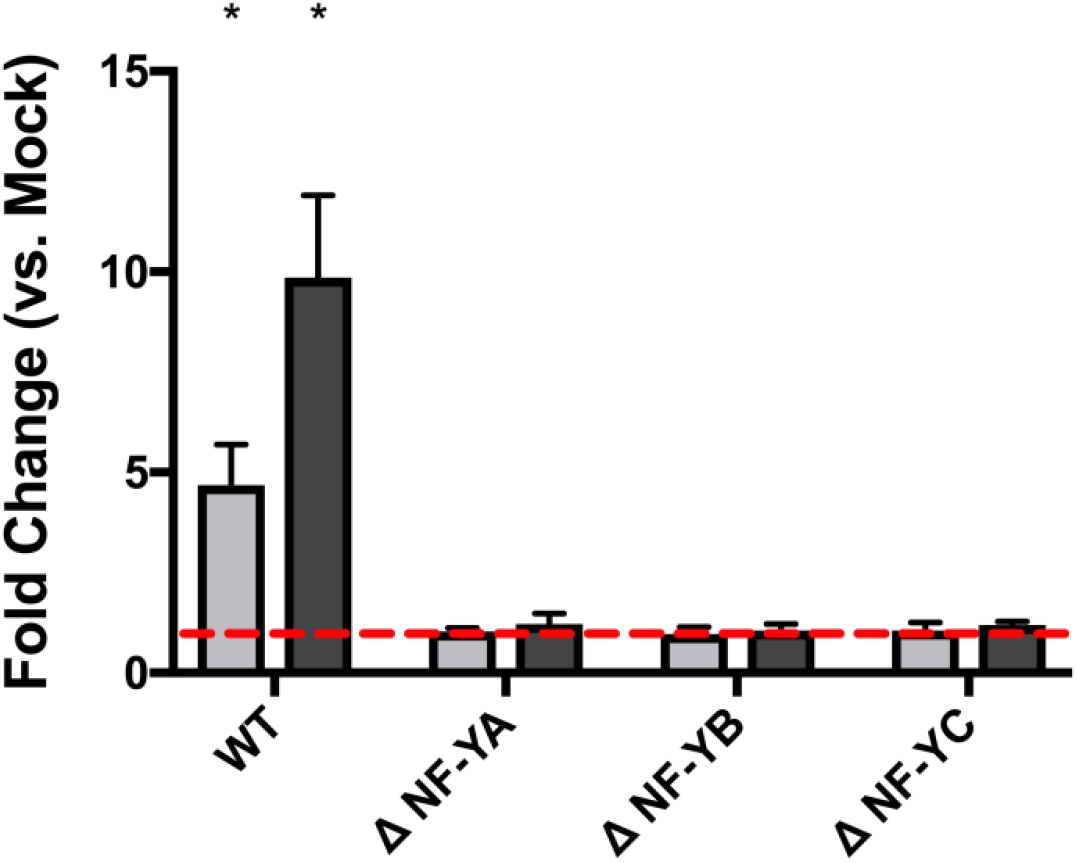
Activation of the DIMR system in the absence of individual NF-Y effectors. NF-^YA2/B2/C3^ Effectors with empty components for NF-YA (ΔNF-YA), NF-YB (ΔNF-YB), or NF-YC (ΔNF-YC), were tested for ability to activate the 4x *FT CCAAT* box in the *ASN1*-based promoter. Error bars represent 95% confidence intervals. Asterisks above individual bars indicate statistical significance over matched mock-treated samples, as determined through two-way ANOVA (p < 0.01); all bars lacking asterisks were found to be nonsignificant compared to matched mock-treated samples.

