## Supplemental Table 1 for "DIMR, a Yeast-Based Synthetic Reporter System for Probing Oligomeric Transcription Factor DNA Binding"

| | | | | |
|---|---|---|---|---|
| pDIMR_A### | Activator Module | | pRS315 backbone | |
| pDIMR_E### | Effector Module | | pRS314 backbone | |
| pDIMR_R### | Reporter Module | | pRSII326 backbone (modified to remove an NcoI site from URA selection) | |
| **Plasmid designation** | **E. coli selection** | **Yeast selection** | **Details** | **pZM designation** |
| pDIMR_A001 | amp | LEU- | pACT1::Z4EV::CYC1b terminator | pZM12 |
| pDIMR_E001 | amp | TRP- | pZ4EV::NF-YA2--3xHA--T2A--NF-YB2--3xMYC--T2A--NF-YC3--3xFLAG::CYC1b terminator | pZM2 |
| pDIMR_E002 | amp | TRP- | pDIMR_E001 with NF-YA2 H183A point mutation | pZM35 |
| pDIMR_E003 | amp | TRP- | pDIMR_E001 with NF-YB2 E65R point mutation | pZM23 |
| pDIMR_E004 | amp | TRP- | pDIMR_E001 with broken T2A site between NF-YA2 and NF-YB2 | pZM51 |
| pDIMR_E005 | amp | TRP- | pDIMR_E001 with broken T2A site between NF-YB2 and NF-YC3 | pZM52 |
| pDIMR_E006 | amp | TRP- | pDIMR_E001 with both T2A sites broken | pZM57 |
| pDIMR_E007 | amp | TRP- | pZ4EV::NF-YB2--3xMYC--T2A--NF-YC3--3xFLAG::CYC1b terminator | pZM61 |
| pDIMR_E008 | amp | TRP- | pZ4EV::NF-YA2--3xHA--T2A--NF-YC3--3xFLAG::CYC1b terminator | pZM106 |
| pDIMR_E009 | amp | TRP- | pZ4EV::NF-YA2--3xHA--T2A--NF-YB2--3xMYC::CYC1b terminator | pZM62 |
| pDIMR_R001 | amp | URA- | 1x FT CCAAT +/- 20bp :: pCIT1 :: Firefly LUC :: CYC1b terminator :: ADH1 terminator :: Renilla LUC :: pADH1 | pZM42 |
| pDIMR_R002 | amp | URA- | 1x FT CCAAT +/- 20bp :: pASN1 :: Firefly LUC :: CYC1b terminator :: ADH1 terminator :: Renilla LUC :: pADH1 | pZM43 |
| pDIMR_R003 | amp | URA- | 1x FT CCAAT +/- 10bp :: pASN1 :: Firefly LUC :: CYC1b terminator :: ADH1 terminator :: Renilla LUC :: pADH1 | pZM53 |
| pDIMR_R004 | amp | URA- | 3x FT CCAAT +/- 10bp :: pASN1 :: Firefly LUC :: CYC1b terminator :: ADH1 terminator :: Renilla LUC :: pADH1 | pZM54 |
| pDIMR_R005 | amp | URA- | 2x FT CCAAT +/- 10bp :: pASN1 :: Firefly LUC :: CYC1b terminator :: ADH1 terminator :: Renilla LUC :: pADH1 | pZM58 |
| pDIMR_R006 | amp | URA- | 2x FT CCAAT +/- 20bp :: pASN1 :: Firefly LUC :: CYC1b terminator :: ADH1 terminator :: Renilla LUC :: pADH1 | pZM59 |
| pDIMR_R007 | amp | URA- | 4x FT CCAAT +/- 20bp :: pASN1 :: Firefly LUC :: CYC1b terminator :: ADH1 terminator :: Renilla LUC :: pADH1 | pZM60 |
| pDIMR_R008 | amp | URA- | pDIMR_R007 CCAAT permutation: CCAAT -> CCAAA | pZM89 |
| pDIMR_R009 | amp | URA- | pDIMR_R007 CCAAT permutation: CCAAT -> CCAAC | pZM90 |
| pDIMR_R010 | amp | URA- | pDIMR_R007 CCAAT permutation: CCAAT -> CCAAG | pZM91 |
| pDIMR_R011 | amp | URA- | pDIMR_R007 CCAAT permutation: CCAAT -> CCACA | pZM92 |
| pDIMR_R012 | amp | URA- | pDIMR_R007 CCAAT permutation: CCAAT -> CCACC | pZM93 |
| pDIMR_R013 | amp | URA- | pDIMR_R007 CCAAT permutation: CCAAT -> CCACG | pZM94 |
| pDIMR_R014 | amp | URA- | pDIMR_R007 CCAAT permutation: CCAAT -> CCACT | pZM95 |
| pDIMR_R015 | amp | URA- | pDIMR_R007 CCAAT permutation: CCAAT -> CCAGA | pZM96 |
| pDIMR_R016 | amp | URA- | pDIMR_R007 CCAAT permutation: CCAAT -> CCAGC | pZM97 |
| pDIMR_R017 | amp | URA- | pDIMR_R007 CCAAT permutation: CCAAT -> CCAGG | pZM98 |
| pDIMR_R018 | amp | URA- | pDIMR_R007 CCAAT permutation: CCAAT -> CCAGT | pZM99 |
| pDIMR_R019 | amp | URA- | pDIMR_R007 CCAAT permutation: CCAAT -> CCATA | pZM100 |
| pDIMR_R020 | amp | URA- | pDIMR_R007 CCAAT permutation: CCAAT -> CCATC | pZM101 |
| pDIMR_R021 | amp | URA- | pDIMR_R007 CCAAT permutation: CCAAT -> CCATG | pZM102 |
| pDIMR_R022 | amp | URA- | pDIMR_R007 CCAAT permutation: CCAAT -> CCATT | pZM103 |
